# Variation-preserving normalization unveils blind spots in gene expression profiling

**DOI:** 10.1101/021212

**Authors:** Carlos P. Roca, Susana I. L. Gomes, Mónica J. B. Amorim, Janeck J. Scott-Fordsmand

**Affiliations:** Department of Chemical Engineering, Universitat Rovira i Virgili, 43007 Tarragona, Spain; Department of Biology & CESAM, University of Aveiro, 3810-193 Aveiro, Portugal; Department of Bioscience, Aarhus University, 8600 Silkeborg, Denmark

**Keywords:** differential gene expression, gene expression microarrays, RNA-Seq, data normalization

## Abstract

RNA-Seq and gene expression microarrays provide comprehensive profiles of gene activity, but lack of reproducibility has hindered their application. A key challenge in the data analysis is the normalization of gene expression levels, which is currently performed following an implicit assumption that most genes are not differentially expressed. Here, we present a mathematical approach to normalization that makes no assumption of this sort. We have found that variation in gene expression is much greater than currently believed, and that it can be measured with available technologies. Our results also explain, at least partially, the problems encountered in transcriptomics studies. We expect this improvement in detection to help efforts to realize the full potential of gene expression profiling, especially in analyses of cellular processes involving complex modulations of gene expression.

## Introduction

Since the discovery of DNA structure by Watson and Crick, molecular biology has progressed increasingly quickly, with rapid advances in sequencing and related genomic technologies. Among these, microarrays and RNA-Seq have been widely adopted to obtain gene expression profiles, by measuring the concentration of tens of thousands of mRNA molecules in single assays (Schena et al., 1995; Lockhart et al., 1996; Duggan et al., 1999; Mortazavi et al., 2008; Wang et al., 2009). Despite their enormous potential (Golub et al., 1999; van ’t Veer et al., 2002; Ivanova et al., 2002; Chi et al., 2003), problems of reproducibility and reliability (Tan et al., 2003; Frantz, 2005; Couzin, 2006) have discouraged their use in some areas, e.g. biomedicine (Michiels et al., 2005; Weigelt and Reis-Filho, 2010; Brettingham-Moore et al., 2011). In more mature microarray technologies, issues such as probe design, cross-hybridization, non-linearities and batch effects (Draghici et al., 2006) have been identified as possible culprits, but the problems persist (Shi et al., 2006; Su et al., 2014).

The normalization of gene expression, which is required to set a common reference level among samples (Smyth and Speed, 2003; Irizarry et al., 2003; Bullard et al., 2010; Garber et al., 2011; Dillies et al., 2013), is also reportedly problematic, affecting the reproducibility of results with both microarray (Shi et al., 2006; Shippy et al., 2006) and RNA-Seq (Su et al., 2014; Bullard et al., 2010; Dillies et al., 2013). Batch effects and their influence on normalization have recently received a great deal of attention (Leek et al., 2010; Reese et al., 2013; Li et al., 2014), resulting in approaches aiming to remove unwanted technical variation caused by differences between batches of samples or by other sources of expression heterogeneity (Listgarten et al., 2010; Gagnon-Bartsch and Speed, 2012; Risso et al., 2014). A different issue, however, is the underlying assumption made by the most widely used normalization methods to date, such as median and quantile normalization (Bolstad et al., 2003) for microarrays, or RPKM (Mortazavi et al., 2008) and TMM (Robinson and Oshlack, 2010) for RNA-Seq, which posit that most genes are not differentially expressed (Dillies et al., 2013; Hicks and Irizarry, 2015). This lack-of-variatio*n assumption ma*y seem reasonable for many applications, but it has not been confirmed. Furthermore, results obtained with other technologies, particularly qRT-PCR, suggest that it may not be valid (Shi et al., 2006; Bullard et al., 2010).

Some methods have been proposed to address the issue of the lack-of-variation assumption, based on the use of spike-ins (Lovén et al., 2012), negative control probes (Wu and Aryee, 2010) or negative control genes (Gagnon-Bartsch and Speed, 2012), that is, on external or internal controls that are *known a priori* not to be differentially expressed (Lippa et al., 2010). The applicability of these methods, however, has been limited by this requirement of a priori knowledge, which is rarely available for a sufficiently large number of controls. Thus, in attempts to clarify and overcome limitations imposed by the lack-of-variation assumption, we have developed an approach to normalization that does not assume lack-of-variation and that does not require the use of spike-ins or a priori knowledge of control genes. The analysis of a large gene expression dataset using this approach shows that the assumption can severely undermine the detection of variation in gene expression. We have found that large numbers of differentially expressed genes with substantial expression changes are missed when data are normalized with methods that assume lack-of-variation.

## Results

### Datasets and Normalization Methods

The dataset was obtained from biological triplicates of *Enchytraeus crypticus* (a globally distributed soil organism used in standard ecotoxicity tests), sampled under 51 experimental conditions (42 treatments and 9 controls), involving exposure to several substances, at several concentrations and durations according to a factorial design (Supp. Table 1). Gene expression was measured using a customized high-density oligonucleotide microarray, and the resulting dataset was normalized with four methods. Two of these methods are the most widely used procedures for microarrays, median (or scale) normalization and quantile normalization (Bolstad et al., 2003), whereas the other two, designated *median condition-decomposition normalization* and *standard-vector condition-decomposition normalization*, have been developed for this study.

With the exception of quantile normalization, all used methods apply a multiplicative factor to the expression levels in each sample, equivalent to the addition of a number in the usual log_2_-scale for gene expression levels. Solving the *normalization problem* consists of finding these correction factors. The problem can be exactly and linearly decomposed into several sub-problems: one within-condition normalization for each experimental condition and one final between-condition normalization for the condition averages. In the within-condition normalizations, the samples (replicates) subjected to each experimental condition are normalized separately, whereas in the final between-condition normalization average levels for all conditions are normalized together. Because there are no genes with differential expression in any of the within-condition normalizations, the lack-of-variation assumption only affects the final between-condition normalization. The assumption is avoided by using, in this normalization, expression levels only from *no-variation genes*, i.e. genes that show no evidence of differential expression under a statistical test. Both methods of normalization proposed here follow this condition-decomposition approach.

With median condition-decomposition normalization, all normalizations are performed are included in the between-condition step. Otherwise, if all genes were used in this final step, the resulting total normalization factors would be exactly the same as those obtained with conventional median normalization.

For standard-vector condition-decomposition normalization, a vectorial procedure was developed to carry out each normalization step. The samples of any experimental condition, in a properly normalized dataset, must be *exchangeable*. In mathematical terms, the expression levels of each gene can be considered as an *s*-dimensional vector, where *s* is the number of samples for the experimental condition. After standardization (mean subtraction and variance scaling), these standard vectors are located in a (*s* − 2)-dimensional hypersphere. The exchangeability mentioned above implies that, when properly normalized, the distribution of standard vectors must be invariant with respect to permutations of the sample labels and must have zero expected value. These properties allow to obtain, under fairly general assumptions, a robust estimator of the normalization factors.

To further explore and compare outcomes of the normalization methods, they were also applied to a synthetic random dataset. This dataset was generated with identical means and variances gene-by-gene to the real dataset, and with the assumption that all genes were no-variation genes. In addition, normalization factors were applied, equal to those obtained from the real dataset. Thus, the synthetic dataset was very similar to the real one, while complying by construction with the lack-of-variation assumption.

### Normalization Results

Figure 1 displays the results of applying the four normalization methods to the real and synthetic datasets. Each panel shows the interquartile ranges of expression levels for the 153 samples, grouped in triplicates exposed to each experimental condition. Both median (second row) and quantile normalization (third row) yielded similar outputs, for both datasets. In contrast, the condition-decomposition normalizations (fourth and fifth rows) identified marked differences, detecting much greater variation between conditions in the sample the same, while quantile normalization makes the full distribution of each sample the same. Hence, if there were differences in medians or distributions of gene expression between experimental conditions, both methods would have removed them. Figures 1G,I show that such variation between conditions was present in the real dataset.

**Figure 1:**
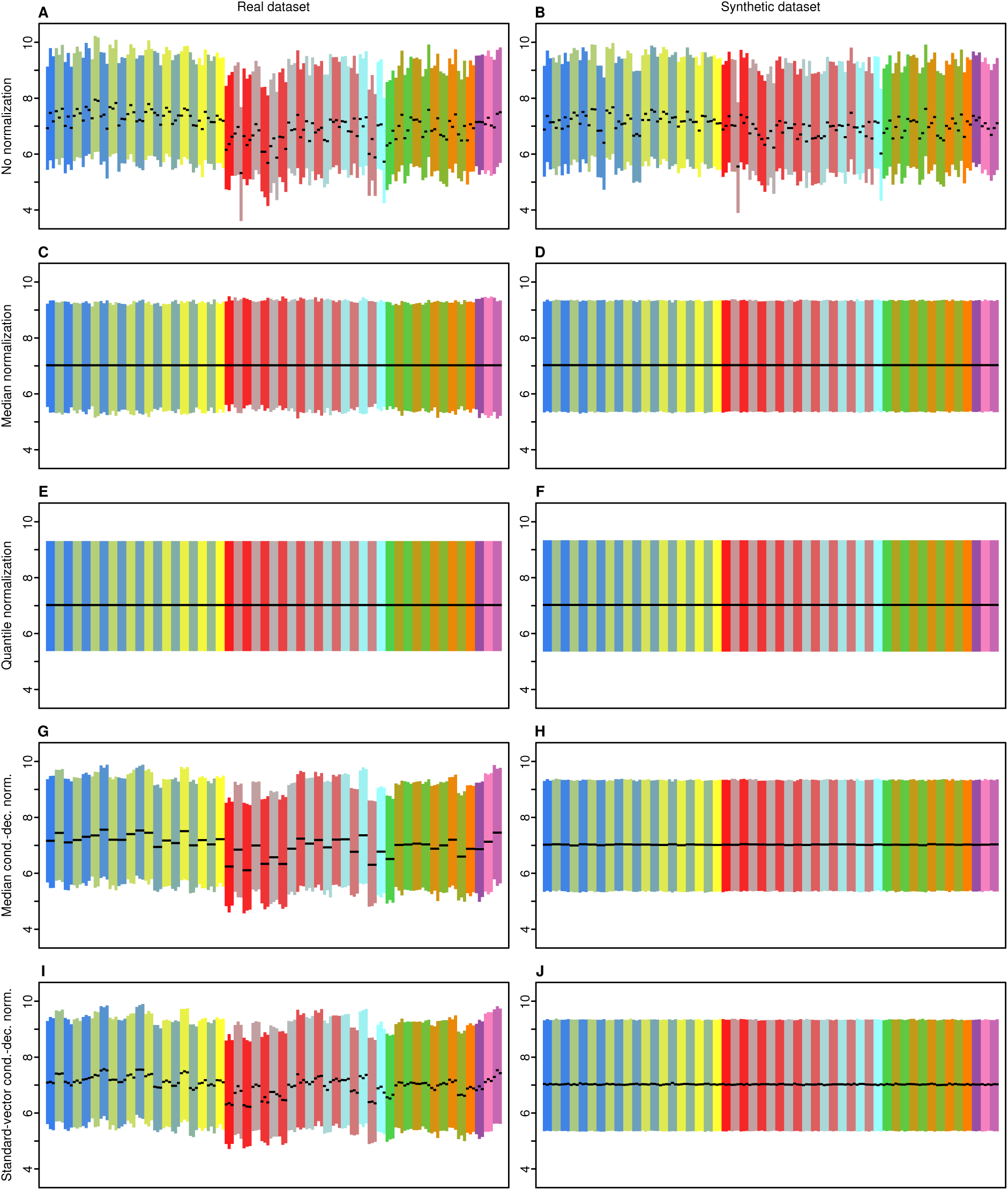
The condition-decomposition normalizations detected a large amount of between-condition variation in the real expression data, in contrast with conventional methods. All 10 panels show interquartile ranges of expression levels of the 153 samples, grouped by the 51 experimental conditions (Ag, blue-yellow; Cu, red-cyan; Ni, green-orange; UV, purple; see Supp. Table 1). Black lines indicate medians. Rows and columns correspond to normalization methods and datasets (as labeled), respectively. In the synthetic dataset no gene was differentially expressed between any two conditions.

### Influence of no-variation genes on normalization

To clarify how the condition-decomposition normalizations preserved the variation between conditions, we studied the influence of the choice of no-variation genes in the final between-condition normalization. To this end, we obtained the between-condition variation with both methods in two families of cases. In one family, no-variation genes were chosen in decreasing order of *p*-values from an ANOVA test. In the other family, genes were chosen at random. The first option was similar to the approach implemented to obtain the results presented in Figures 1G–J, with the difference that there the number of genes was chosen automatically by a statistical test. As shown in Figure 2A, for the real dataset the random choice of genes resulted in *n*^₋1/2^ decays (*n* being the number of chosen genes), followed by a plateau. The *n*^₋1/2^ decays reflect the standard errors of the estimators of the normalization factors. Selecting the genes by decreasing *p*-values, however, yielded a completely different result. Up to a certain number of genes, the variance remained similar, but for larger numbers of genes the variance dropped rapidly. Figure 2A shows, therefore, that between-condition variation was removed as soon as the between-condition normalizations used genes that varied in expression level across experimental conditions. The big circles in Figure 2A indicate the working points of the normalizations used to generate the results displayed in Figures 1G,I. In fact, these points slightly underestimated the variation between conditions. Although the statistical test for identifying no-variation genes ensured that there was no evidence of variation, inevitably the expression of some selected genes varied across conditions.

**Figure 2:**
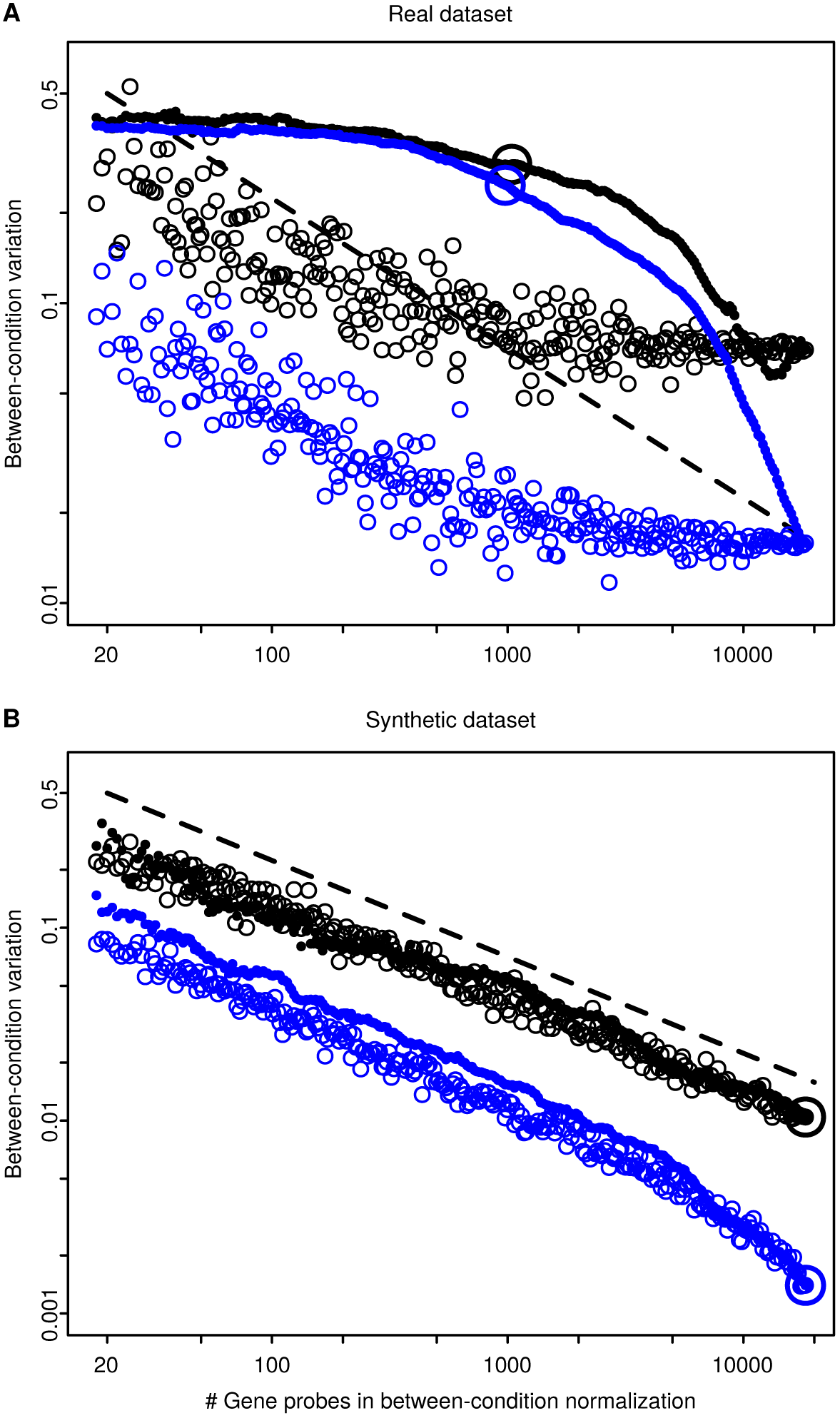
The selection of genes in the final between-condition normalization was crucial to preserve variation between conditions. The panels show the detected variation as a function of the number of gene probes used in the between-condition normalization of the real dataset (A) and synthetic dataset (B). Between-condition variation is represented as the standard deviation of the within-condition mean averages (averages of sample mean expression levels, for all samples under the condition). See Supplementary for within-condition median averages, with similar results. Each point in either of the panels indicates the variation obtained with one complete normalization (black circles, median condition-decomposition normalization; blue circles, standard-vector condition-decomposition normalization). Gene probes were selected in two ways: randomly (empty circles) or in decreasing order of *p*-values (filled circles). Big circles show the working points of the algorithms whose results are depicted in Figures 1G–J. Black dashed lines show references for *n*^−1/2^ decays, with the same values in both panels.

Figure 2B displays the results obtained with the synthetic dataset. There were no plateaus when no-variation genes were chosen randomly, only *n*^₋1/2^ decays, and small differences when no-variation genes were selected by decreasing *p*-values. Big circles show that working points were selected with much larger numbers of genes in the synthetic dataset (Figs. 1H,J) than in the real dataset (Figs. 1G,I). The residual variation, produced by errors in the estimation of the normalization factors, was much smaller than the variation detected in the real dataset, especially for standard-vector condition-decomposition normalization. Overall, Figure 2 shows that the between-condition variation pictured in Figures 1G,I is not an artifact caused by using an exceedingly small or extremely particular set of genes in the final between-condition normalization, but that this variation originated from the real dataset.

### Differential Gene Expression

Finally, Figure 3A shows the numbers of differentially expressed gene probes (DEGP), identified after normalizing with the four methods, for each of the 42 experimental treatments versus the corresponding control (Supp. Table 2). Compared to conventional methods, the number of DEGP detected with the condition-decomposition normalizations was much larger under most treatments, including some whose number of DEGP was larger by more than one order of magnitude. These are statistically significant changes of gene expression, i.e. changes that cannot be explained by chance. More important is the scale of the detected variation, as illustrated by the boxplots in Figure 3C showing absolute fold changes of DEGP detected after standard-vector condition-decomposition normalization. For all treatments, the entire interquartile range of absolute fold change is above 1.5-fold, and for more than two thirds of the treatments the median absolute fold change is greater than 2. This amount of gene expression variation cannot be neglected, and warrants further research to explore its biological significance.

**Figure 3:**
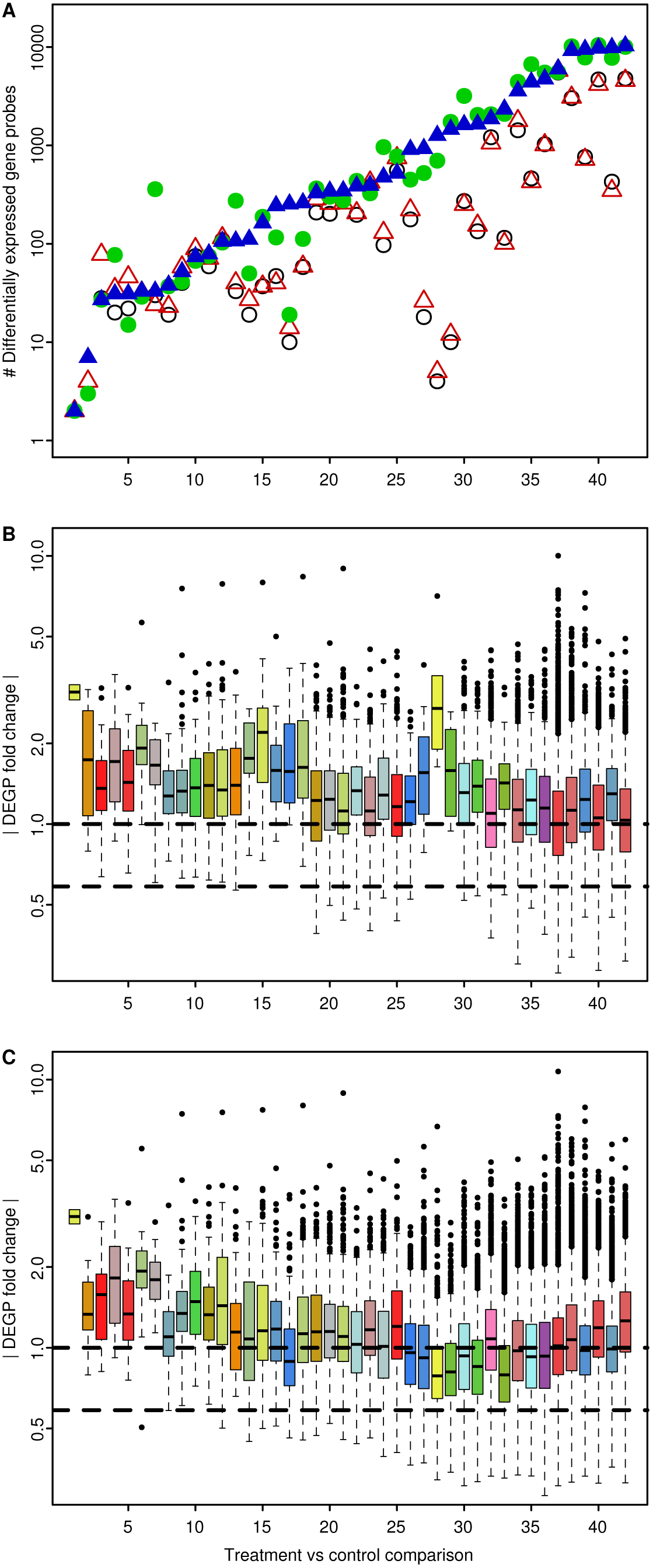
The condition-decomposition normalizations detected much larger numbers of differentially expressed gene probes (DEGP), with substantial fold changes. A: Number of DEGP for each treatment vs control comparison, obtained after applying the four normalization methods (empty black circles, median normalization; empty red triangles, quantile normalization; filled green circles, median condition-decomposition normalization; filled blue triangles, standard-vector condition decomposition normalization). Significant differential expression was identified with R/Bioconductor package limma. (see Supp. Fig. 2 for results with t-tests). Lower panel shows boxplots of absolute values of DEGP fold changes (absolute differences of log_2_ expression levels), also per treatment vs control comparison, obtained with quantile normalization (B) and standard-vector condition-decomposition normalization (C). Boxplots are colored by treatment, with the same color code as in . In both panels comparisons are ordered according to the number of DEGP identified with standard-vector condition-decomposition normalization, increasing from left to right (Supp. Table 2). Dashed horizontal lines in the lower panel indicate references of 1.5-fold and 2-fold changes.

## Discussion

The variation between medians displayed in Figures 1G,I may seem surprising, given routine expectations based on current methods (Figs. 1C,E). Nevertheless, this variation inevitably results from the imbalance between over-and under-expressed genes. As an illustration, let us consider a case with two experimental conditions, in which the average expression of a given gene is less than the distribution median under one condition, but greater than the median under the other. The variation of this gene alone will change the value of the median to the expression level of the next ranked gene. Therefore, if the number of over-expressed genes is different from the number of under-expressed genes, and enough changes cross the median boundary, then the median will substantially differ between conditions. Only when the differential expression is balanced or small enough, will the median stay the same. This argument applies equally to any other quantile in the distribution of gene expression. Transcriptional amplification is an extreme example of change in the distribution of expression levels (Lovén et al., 2012), which can nevertheless be properly normalized with condition-decomposition methods, and without resorting to spike-ins as long as some genes are not differentially expressed.

An important feature of the approaches to normalization proposed here (linear decomposition into normalization sub-problems per condition, and standard-vector normalization for each sub-problem) is that they do not depend on any particular aspect of the technology of gene expression microarrays or RNA-Seq. The numbers in the input data are interpreted as measured concentrations of mRNA molecules, in order to identify the normalization factors and irrespectively of whether the concentrations were obtained from fluorescence intensities of hybridized cDNA (microarrays) or from counts of fragments read of mRNA sequences (RNA-Seq). Nevertheless, we consider that specific within-sample corrections for each technology are still necessary and must be applied *before* the between-sample normalizations proposed here. Examples include background correction for microarrays or gene-length normalization (RPKM) for RNA-Seq. Equally, methods that address the influence of biological or technical confounding factors on downstream applied when necessary, *after* normalizing.

The lack-of-variation assumption underlying the current methods of normalization was self-fulfilling, removing variation in gene expression that was present in the real dataset. Moreover, it had negative consequences for downstream analyses, as it both removed potentially important biological information and introduced errors in the detection of gene expression. A removal of variation can be understood as errors in the estimation of normalization factors. Considering data and errors vectorially, the length of each vector equals, after centering and up to a constant factor, the standard deviation of the data or error. The addition of an error of small magnitude, compared to the data variance, would have only a minor effect. However, errors of similar or greater magnitude than the data variance may, depending on the lengths and relative angles of the vectors, severely distort the observed data variance. This will in turn cause spurious results in the statistical analyses. Furthermore, the angles between the data and the correct normalization factors (considered as vectors) are random. Data reflect biological variation, while normalization factors respond to technical variation. If the experiment is repeated, even with exactly the same experimental settings, the errors in the normalization factors will vary randomly, causing random spurious results in the downstream analyses. This explains, at least partially, the lack of reproducibility found in transcriptomics studies, especially for the detection of small changes of gene expression, because small variations are most likely to be distorted by errors in the estimates of normalization factors. Accordingly, the largest differences in numbers of DEGP detected by conventional compared to condition-decomposition methods (Fig. 3A) occurred consistently in the treatments with the smallest magnitudes of gene expression changes, e.g. treatments 28, 29 and 33 (Figs. 3B,C).

In summary, this study proves that large numbers of genes change in expression level (often strongly) across experimental conditions, and too extensively to ignore in the normalization of gene expression data. Further, our approach, which avoids the prevailing lack-of-variation assumption, demonstrates that current normalization methods likely remove and distort important variation in gene expression. It also offers a means to investhis to provide revealing insights about diverse biomolecular processes, particularly those involving substantial numbers of genes, such as cell differentiation, toxic responses, diseases with non-Mendelian inheritance patterns and cancer. After years of lagging behind the advances in genome sequencing, we believe that the procedures presented here will assist efforts to realize the full potential of gene expression profiling.

## Data Deposition and Code Availability

MIAME-compliant microarray data from the experiment were submitted to the Gene Expression Omnibus (GEO) at the NCBI website (platform: GPL20310; series: GSE69746, GSE69792, GSE69793 and GSE69794). Custom code that reproduces all the reported results starting from the raw microarray data is available at the GitHub repository https://github/carlosproca/gene-expr-norm-paper.

## Acknowledgements

This work was funded by the European Union FP7 projects MODERN (Ref. 309314-2) (C.P.R., J.J.S.-F.) and MARINA (Ref. 263215) (J.J.S.-F.), by FEDER through COMPETE (Programa Operacional Factores de Competitividade) and FCT (Funda¸c˜ao para a Ciˆencia e Tecnologia) through project bio-CHIP (Ref. FCT EXPL/AAG-MAA/0180/2013) (S.I.L.G., M.J.B.A.), and by a post-doctoral grant (Ref. SFRH/BPD/95775/2013) (S.I.L.G).

## Author Contributions

S.I.L.G., M.J.B.A. and J.J.S.-F. designed the toxicity experiment. S.I.L.G. carried out the experimental work and collected the microarray data. C.P.R. designed and implemented the novel normalization methods. C.P.R. performed the statistical analyses. All the authors jointly discussed the results. C.P.R. drafted the paper, with input from all the authors. All the authors edited the final version of the paper.

## Materials and Methods

### Test Organism and Exposure Media

The test species was *Enchytraeus crypticus*. Individuals were cultured in Petri dishes containing agar medium, in controlled conditions (Gomes et al., 2015b).

For copper (Cu) exposure, a natural soil collected at Hygum, Jutland, Denmark was used (Gomes et al., 2015b; Scott-Fordsmand et al., 2000). For silver (Ag) and nickel (Ni) exposure, the natural standard soil LUFA 2.2 (LUFA Speyer, Germany) was used (Gomes et al., 2015b). The exposure to ultra-violet (UV) radiation was done in ISO reconstituted water (OECD, 2004a).

### Test Chemicals

The tested Cu forms (Gomes et al., 2015b) included copper nitrate (Cu(NO3)2 ·3H2O *>* 99%, Sigma Aldrich), Cu nanoparticles (Cu-NPs, 20–30 nm, American Elements) and Cu nanowires (Cu-Nwires, synthesized by reduction of copper (II) nitrate with hydrazine in alkaline medium (Chang et al., 2005)).

The tested Ag forms (Gomes et al., 2015b) included silver nitratre AgNO_3_ *>* 99%, Sigma Aldrich), non-coated Ag nanoparticles (Ag-NPs Non-Coated, 20–30 nm, American Elements),

Polyvinylpyrrolidone (PVP)-coated Ag nanoparticles (Ag-NPs PVP-Coated, 20–30 nm, American Elements), and Ag NM300K nanoparticles (Ag NM300K, 15 nm, JRC Repository). The Ag NM300K was dispersed in 4% Polyoxyethylene Glycerol Triolaete and Polyoxyethylene (20) orbitan mono-Laurat (Tween 20), thus the dispersant was tested alone as control (CTdisp).

The tested Ni forms included nickel nitrate (Ni(NO_3_)_2_ ·6H2O ≥ 98.5%, Fluka) and Ni nanoparticles (Ni-NPs, 20 nm, American Elements).

### Spiking Procedure

Spiking for the Cu and Ag materials was done as in previous work (Gomes et al., 2015b). For the Ni materials, the Ni-NPs were added to the soil as powder, following the same procedure as for the Cu materials. NiNO_3_, being soluble, was added to the pre-moistened soil as aqueous dispersions.

The concentrations tested were selected based on the reproduction effect concentrations EC_20_ and EC_50_, for *E. crypticus*, within 95% of confidence intervals, being: CuNO_3_ EC_20/50_ = 290/360 mgCu/kg, Cu-NPs EC_20/50_ = 980/1760 mgCu/kg, Cu-Nwires EC_20/50_ = 850/1610 mgCu/kg, Cu-Field EC_20/50_ = 500/1400 mgCu/kg, AgNO_3_ EC_20/50_ = 45/60 mgAg/kg, Ag-NP PVP-coated EC_20/50_ = 380/550 mgAg/kg, Ag-NP Non-coated EC_20/50_ = 380/430 mgAg/kg, Ag NM300K EC_20/50_ = 60/170 mgAg/kg, CTdisp = 4% w/w Tween 20, NiNO_3_ EC_20/50_ = 40/60 mgNi/kg, Ni-NPs EC_20/50_ = 980/1760 mgNi/kg.

Four biological replicates were performed per test condition, including controls. For Cu exposure, the control condition for all the treatments consisted of soil from a control area at Hygum site, which has a Cu background concentration of 15 mg/kg (Scott-Fordsmand et al., 2000). For Ag exposure, two control sets were performed: CT (un-spiked LUFA soil, to be the control condition for AgNO_3_, Ag-NPs PVP-Coated and Ag-NPs Non-Coated treatments) and CTdisp (LUFA soil spiked with the dispersant Tween 20, to be the control condition for the Ag NM300K treatments). For Ni exposure, the control consisted of un-spiked LUFA soil.

### Exposure Details

In soil (i.e. for Cu, Ag and Ni) exposure followed the standard ERT (OECD, 2004b) with adaptations as follows: twenty adults with well-developed clitellum were introduced in each test vessel, containing 20 g of moist soil (control or spiked). The organisms were exposed for three and seven days under controlled conditions of photoperiod (16:8 h light:dark) and temperature 20 ± 1 ^◦^C without food. After the exposure period, the liquid nitrogen. The samples were stored at −80 ^◦^C, until analysis.

For UV exposure, the test conditions (OECD, 2004a) were adapted for *E. crypticus* (Gomes et al., 2015a). The exposure was performed in 24-well plates, where each well correspond to a replicate and contain 1 ml of ISO water and five adult organisms with clitellum. The test duration was five days, at 20 ± 1 ^◦^C. The organisms were exposed to UV on a daily basis, during 15 minutes per day to two UV intensities (280–400nm) of 1669.25 ± 50.83 and 1804.08 ± 43.10 mW*/*m^2^, corresponding to total UV doses of 7511.6 and 8118.35 J*/*m^2^, respectively. The remaining time was spent under standard laboratory illumination (16:8 h photoperiod). UV radiation was provided by an UV lamp (Spectroline XX15F/B, Spectronics Corporation, NY, USA, peak emission at 312 nm) and a cellulose acetate sheet was coupled to the lamp to cut-off UVC-range wavelengths (Gomes et al., 2015a). Thirty two replicates per test condition (including control without UV radiation) were performed to obtain 4 biological replicates with 40 organisms each for RNA extraction. After the exposure period, the organisms were carefully removed from the water and frozen in liquid nitrogen. The samples were stored at −80 ^◦^C, until analysis.

### RNA Extraction, Labeling and Hybridization

RNA was extracted from each replicate, which contained a pool of 20 and 40 organisms, for soil and water exposure, respectively. Three biological replicates per test treatment (including controls) were used. Total RNA was extracted using SV Total RNA Isolation System (Promega). The quantity and purity were measured spectrophotometrically with a nanodrop (NanoDrop ND-1000 Spectrophotometer) and its quality checked by denaturing formaldehyde agarose gel electrophoresis.

500 ng of total RNA were amplified and labeled with Agilent Low Input Quick Amp Labeling Kit (Agilent Technologies, Palo Alto, CA, USA). Positive controls were added with the Agilent one-color RNA Spike-In Kit. Purification of the amplified and labeled cRNA was performed with RNeasy columns (Qiagen, Valencia, CA, USA).

The cRNA samples were hybridized on custom Gene Expression Agilent Microarrays (4 x 44k format), with a single-color design (Castro-Ferreira et al., 2014). Hybridizations were performed using the Agilent Gene Expression Hybridization Kit and each biological replicate was individually hybridized on one array. The arrays were hybridized at 65 ^◦^C with a rotation of 10 rpm, during 17 h. Afterwards, microarrays were washed using Agilent Gene Expression Wash Buffer Kit and scanned with the Agilent DNA microarray scanner G2505B.

### Data Acquisition and Analysis

Fluorescence intensity data was obtained with Agilent Feature Extraction Software v. 10.7.3.1, using recommended protocol GE1_107_Sep09. Quality control was done by inspecting the reports on the Agilent Spike-in control probes. Background correction was provided by Agilent Feature Extraction software. To ensure an optimal comparison between the different normalization methods, only gene probes with good signal quality (flag IsPosAndSignif = True) in all samples were employed in the analyses. This implied the selection of 18,339 gene probes from a total of 43,750. Analyses were performed with R (R Core Team, 2015) v. 3.2.2, using R packages plotrix and RColorBrewer, and with Bioconductor (Huber et al., 2015) v. 3.1 packages genefilter and limma (Ritchie et al., 2015).

The synthetic data was generated gene by gene as normal variates with mean and variance equal, respectively, to the sample mean and sample variance of the real data. The applied normalization factors were those detected from the real data with standard-vector condition-decomposition normalization.

Median normalization was performed by subtracting the median of each sample distribution, and then adding the overall median to preserve the global expression level. Quantile normalization was performed as implemented in the limma package.

The two condition-decomposition normalizations proceeded in the same way: first, 51 independent within-condition normalization using all genes; then, final between-condition normalization, iteratively detecting no-variation genes and normalizing until convergence.

No-variation genes were identified with one-sided Kolmogorov-Smirnov tests, as goodness-of-fit tests against the uniform distribution, carried out on the greatest *p*-values obtained from an ANOVA test on the complete dataset (see below). The ANOVA test benefited from the already corrected within-condition variances, provided by the within-condition normalizations. The KS test was rejected at *α* = 0.001.

The criterion for convergence for the median condition-decomposition (CD) normalizations was to require that the relative changes in the standard deviation of the normalization factors were less than 1%, or less than 10% for 10 steps in a row. In the case of standard-vector CD normalizations, convergence required that numerical errors were, compared to the estimated statistical errors (see below), less than 1%, or less than 10% for 10 steps in a row. For Figure 2 and Supplementary Figure 1, due to the very low number of gene probes in some cases, the thresholds for convergence for 10 steps in a row were increased to 80% and 50%, respectively, for median CD and standard-vector CD normalization.

In standard-vector CD normalization, the distribution of standard vectors was trimmed in each step to remove the 1% more extreme values of variance.

Differentially expressed gene probes were identified with limma (Fig. 3) or t-tests (Supp. Fig. 2), using in all cases a FDR threshold of 5%.

The reference distribution with permutation symmetry shown in the polar plots of the probability density function in Supplementary Movies 1–3 was calculated with the 6 permutations of the empirical standard vectors. The Watson *U*^2^ statistic was calculated with the two-sample test (Durbin, 1973). An equal number of samples for comparison was obtained by sampling with replacement the permuted standard vectors.

### Mathematical Methods

In a gene expression dataset with *g* genes, *c* experimental conditions and *n* samples per condition, the *observed* expression levels of gene *j* in condition *k*, 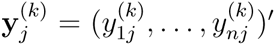, can be expressed in log_2_-scale as

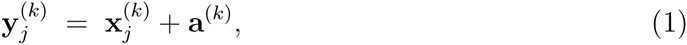
where 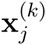 is the vector of *true* gene expression levels and **a**^(k)^ is the vector of normalization factors.

Given a sample vector **x**, the mean vector is 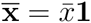, and the residual vector is 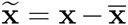.

Then, (1) can be linearly decomposed into

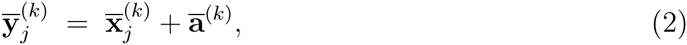

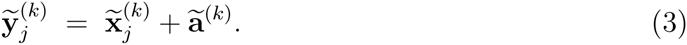

Equations (3) define the within-condition normalizations for each condition *k*. The scalar values in (2) are used to obtain the equations on condition means,

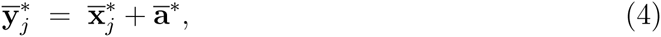

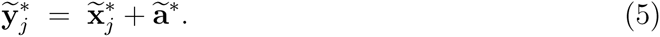

The between-condition normalization is defined by (5). Equations (4) reduce to a single number, which is irrelevant to the normalization. The complete solution for each condition is obtained with 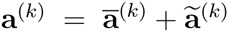.

The *n* samples of gene *j* in a given condition can be modeled with the random vectors 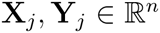. Again, **Y**j = **X***j* + **a**, where **a** is a fixed vector of normalization factors. It can be proved, under fairly general assumptions, that the true standard vectors have zero expected value

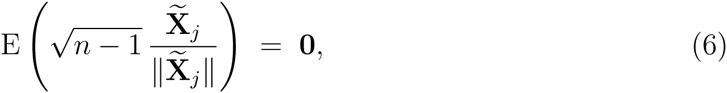

whereas the observed standard vectors verify, as long as **a** ≠ 0,

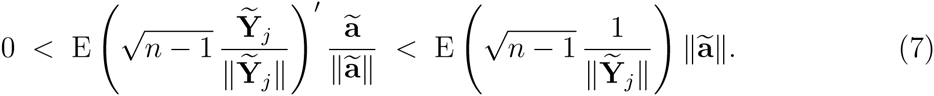

This motivates the following iterative procedure to solve (3) and (5) (*standard-vector normalization*):

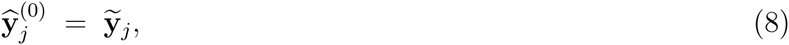

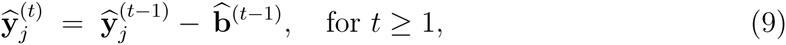

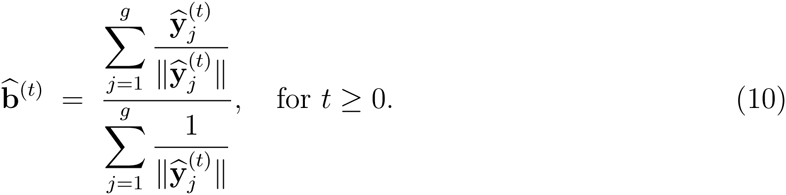

At convergence, 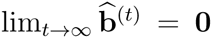, which implies 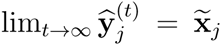 and 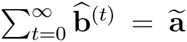. Convergence is faster the more symmetric the empirical distribution of 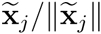 is on the unit (*n* − 2)-sphere. Convergence is optimal with spherically symmetric distributions, such as the Gaussian distribution, because in that case

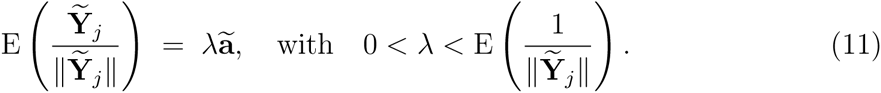

Assuming no correlation between genes, an approximation of the statistical error at step *t* can be obtained with

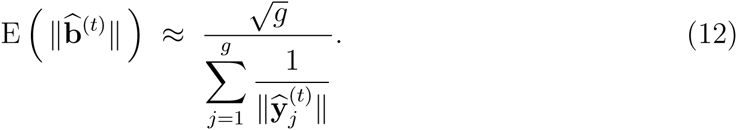

This statistical error is compared with the numerical error to assess convergence.

See Supplementary Material for a detailed exposition of the mathematical methods, and Supplementary Movies 1–5 for an illustration.

## Supplementary Tables

**Supplementary Table 1:** Experimental conditions of the toxicity experiment on *E. crypticus*, listed in the same order as they appear in each panel of, from left to right.

| Condition number | Condition ID | Condition description |
| --- | --- | --- |
| 1 | Ag.AgNO3.EC20.3d | AgNO3 EC20 3 days |
| 2 | Ag.AgNO3.EC20.7d | AgNO3 EC20 7 days |
| 3 | Ag.AgNO3.EC50.3d | AgNO3 EC50 3 days |
| 4 | Ag.AgNO3.EC50.7d | AgNO3 EC50 7 days |
| 5 | Ag.Coated.EC20.3d | Ag-NPs PVP-Coated EC20 3 days |
| 6 | Ag.Coated.EC20.7d | Ag-NPs PVP-Coated EC20 7 days |
| 7 | Ag.Coated.EC50.3d | Ag-NPs PVP-Coated EC50 3 days |
| 8 | Ag.Coated.EC50.7d | Ag-NPs PVP-Coated EC50 7 days |
| 9 | Ag.NC.EC20.3d | Ag-NPs Non-Coated EC20 3 days |
| 10 | Ag.NC.EC20.7d | Ag-NPs Non-Coated EC20 7 days |
| 11 | Ag.NC.EC50.3d | Ag-NPs Non-Coated EC50 3 days |
| 12 | Ag.NC.EC50.7d | Ag-NPs Non-Coated EC50 7 days |
| 13 | Ag.NM300K.EC20.3d | Ag NM300K EC20 3 days |
| 14 | Ag.NM300K.EC20.7d | Ag NM300K EC20 7 days |
| 15 | Ag.NM300K.EC50.3d | Ag NM300K EC50 3 days |
| 16 | Ag.NM300K.EC50.7d | Ag NM300K EC50 7 days |
| 17 | Ag.CT.3d | Ag Control 3 days |
| 18 | Ag.CT.7d | Ag Control 7 days |
| 19 | Ag.CTD.3d | Ag Control Dispersant 3 days |
| 20 | Ag.CTD.7d | Ag Control Dispersant 7 days |
| 21 | Cu.CuNO3.EC20.3d | CuNO3 EC20 3 days |
| 22 | Cu.CuNO3.EC20.7d | CuNO3 EC20 7 days |
| 23 | Cu.CuNO3.EC50.3d | CuNO3 EC50 3 days |
| 24 | Cu.CuNO3.EC50.7d | CuNO3 EC50 7 days |
| 25 | Cu.Cu.NPs.EC20.3d | Cu-NPs EC20 3 days |
| 26 | Cu.Cu.NPs.EC20.7d | Cu-NPs EC20 7 days |
| 27 | Cu.Cu.NPs.EC50.3d | Cu-NPs EC50 3 days |
| 28 | Cu.Cu.NPs.EC50.7d | Cu-NPs EC50 7 days |
| 29 | Cu.Cu.Nwires.EC20.3d | Cu-NWires EC20 3 days |
| 30 | Cu.Cu.Nwires.EC20.7d | Cu-NWires EC20 7 days |
| 31 | Cu.Cu.Nwires.EC50.3d | Cu-NWires EC50 3 days |
| 32 | Cu.Cu.Nwires.EC50.7d | Cu-NWires EC50 7 days |
| 33 | Cu.Cu.field.EC20.3d | Cu-Field EC20 3 days |
| 34 | Cu.Cu.field.EC20.7d | Cu-Field EC20 7 days |
| 35 | Cu.Cu.field.EC50.3d | Cu-Field EC50 3 days |
| 36 | Cu.Cu.field.EC50.7d | Cu-Field EC50 7 days |
| 37 | Cu.CT.3d | Cu Control 3 days |
| 38 | Cu.CT.7d | Cu Control 7 days |
| 39 | Ni.NiNO3.EC20.3d | NiNO3 EC20 3 days |
| 40 | Ni.NiNO3.EC20.7d | NiNO3 EC20 7 days |
| 41 | Ni.NiNO3.EC50.3d | NiNO3 EC50 3 days |
| 42 | Ni.NiNO3.EC50.7d | NiNO3 EC50 7 days |
| 43 | Ni.Ni.NPs.EC20.3d | Ni-NPs EC20 3 days |
| 44 | Ni.Ni.NPs.EC20.7d | Ni-NPs EC20 7 days |
| 45 | Ni.Ni.NPs.EC50.3d | Ni-NPs EC50 3 days |
| 46 | Ni.Ni.NPs.EC50.7d | Ni-NPs EC50 7 days |
| 47 | Ni.CT.3d | Ni Control 3 days |
| 48 | Ni.CT.7d | Ni Control 7 days |
| 49 | Uv.UV.D1.5d | UV Dose 1 |
| 50 | Uv.UV.D2.5d | UV Dose 2 |
| 51 | Uv.CT.5d | UV Control |

**Supplementary Table 2:** Treatment vs control comparisons, listed in increasing number of differentially expressed gene probes (DEGP) obtained with standard-vector condition-decomposition normalization and limma statistical analysis. This is the same order as in, from left to right.

| Comparison number | Treatment ID | Control ID | Treatment description | Number of DEGP |
| --- | --- | --- | --- | --- |
| 1 | Ag.NM300K.EC20.7d | Ag.CTD.7d | Ag NM300K EC20 7 days | 2 |
| 2 | Ni.Ni.NPs.EC20.7d | Ni.CT.7d | Ni-NPs EC20 7 days | 7 |
| 3 | Cu.Cu.NO3.EC20.3d | Cu.CT.3d | CuNO3 EC20 3 days | 27 |
| 4 | Cu.Cu.NO3.EC50.7d | Cu.CT.7d | CuNO3 EC50 7 days | 31 |
| 5 | Cu.Cu.NPs.EC20.3d | Cu.CT.3d | Cu-NPs EC20 3 days | 31 |
| 6 | Ag.AgNO3.EC50.7d | Ag.CT.7d | AgNO3 EC50 7 days | 33 |
| 7 | Cu.Cu.NPs.EC20.7d | Cu.CT.7d | Cu-NPs EC20 7 days | 33 |
| 8 | Ag.NM300K.EC20.3d | Ag.CTD.3d | Ag NM300K EC20 3 days | 38 |
| 9 | Ag.NM300K.EC50.3d | Ag.CTD.3d | Ag NM300K EC50 3 days | 52 |
| 10 | Ni.NiNO3.EC20.3d | Ni.CT.3d | NiNO3 EC20 3 days | 74 |
| 11 | Ni.NiNO3.EC20.7d | Ni.CT.7d | NiNO3 EC20 7 days | 79 |
| 12 | Ag.NC.EC20.7d | Ag.CT.7d | Ag-NPs Non-Coated EC20 7 days | 106 |
| 13 | Ni.Ni.NPs.EC50.7d | Ni.CT.7d | Ni-NPs EC50 7 days | 107 |
| 14 | Ag.AgNO3.EC20.7d | Ag.CT.7d | AgNO3 EC20 7 days | 111 |
| 15 | Ag.NC.EC50.7d | Ag.CT.7d | Ag-NPs Non-Coated EC50 7 days | 163 |
| 16 | Ag.NC.EC20.3d | Ag.CT.3d | Ag-NPs Non-Coated EC20 3 days | 244 |
| 17 | Ag.AgNO3.EC50.3d | Ag.CT.3d | AgNO3 EC50 3 days | 254 |
| 18 | Ag.Coated.EC20.7d | Ag.CT.7d | Ag-NPs PVP-Coated EC20 7 days | 261 |
| 19 | Ni.NiNO3.EC50.7d | Ni.CT.7d | NiNO3 EC50 7 days | 329 |
| 20 | Cu.Cu.NPs.EC50.7d | Cu.CT.7d | Cu-NPs EC50 7 days | 343 |
| 21 | Ag.Coated.EC50.7d | Ag.CT.7d | Ag-NPs PVP-Coated EC50 7 days | 346 |
| 22 | Cu.Cu.Nwires.EC50.7d | Cu.CT.7d | Cu-NWires EC50 7 days | 387 |
| 23 | Cu.Cu.NO3.EC20.7d | Cu.CT.7d | CuNO3 EC20 7 days | 393 |
| 24 | Cu.Cu.Nwires.EC20.7d | Cu.CT.7d | Cu-NWires EC20 7 days | 478 |
| 25 | Cu.Cu.NO3.EC50.3d | Cu.CT.3d | CuNO3 EC50 3 days | 522 |
| 26 | Ag.AgNO3.EC20.3d | Ag.CT.3d | AgNO3 EC20 3 days | 911 |
| 27 | Ag.Coated.EC20.3d | Ag.CT.3d | Ag-NPs PVP-Coated EC20 3 days | 930 |
| 28 | Ag.NM300K.EC50.7d | Ag.CTD.7d | Ag NM300K EC50 7 days | 1,264 |
| 29 | Ni.Ni.NPs.EC20.3d | Ni.CT.3d | Ni-NPs EC20 3 days | 1,460 |
| 30 | Cu.Cu.field.EC20.7d | Cu.CT.7d | Cu-Field EC20 7 days | 1,627 |
| 31 | Ni.NiNO3.EC50.3d | Ni.CT.3d | NiNO3 EC50 3 days | 1,649 |
| 32 | Uv.UV.D2.5d | Uv.CT.5d | UV Dose 2 | 1,864 |
| 33 | Ni.Ni.NPs.EC50.3d | Ni.CT.3d | Ni-NPs EC50 3 days | 2,341 |
| 34 | Cu.Cu.field.EC50.3d | Cu.CT.3d | Cu-Field EC50 3 days | 3,578 |
| 35 | Cu.Cu.field.EC50.7d | Cu.CT.7d | Cu-Field EC50 7 days | 4,412 |
| 36 | Uv.UV.D1.5d | Uv.CT.5d | UV Dose 1 | 4,746 |
| 37 | Cu.Cu.NPs.EC50.3d | Cu.CT.3d | Cu-NPs EC50 3 days | 5,993 |
| 38 | Cu.Cu.field.EC20.3d | Cu.CT.3d | Cu-Field EC20 3 days | 9,225 |
| 39 | Ag.Coated.EC50.3d | Ag.CT.3d | Ag-NPs PVP-Coated EC50 3 days | 9,474 |
| 40 | Cu.Cu.Nwires.EC20.3d | Cu.CT.3d | Cu-NWires EC20 3 days | 9,753 |
| 41 | Ag.NC.EC50.3d | Ag.CT.3d | Ag-NPs Non-Coated EC50 3 days | 9,876 |
| 42 | Cu.Cu.Nwires.EC50.3d | Cu.CT.3d | Cu-NWires EC50 3 days | 10,287 |

## Supplementary Figures

**Supplementary Figure 1:**
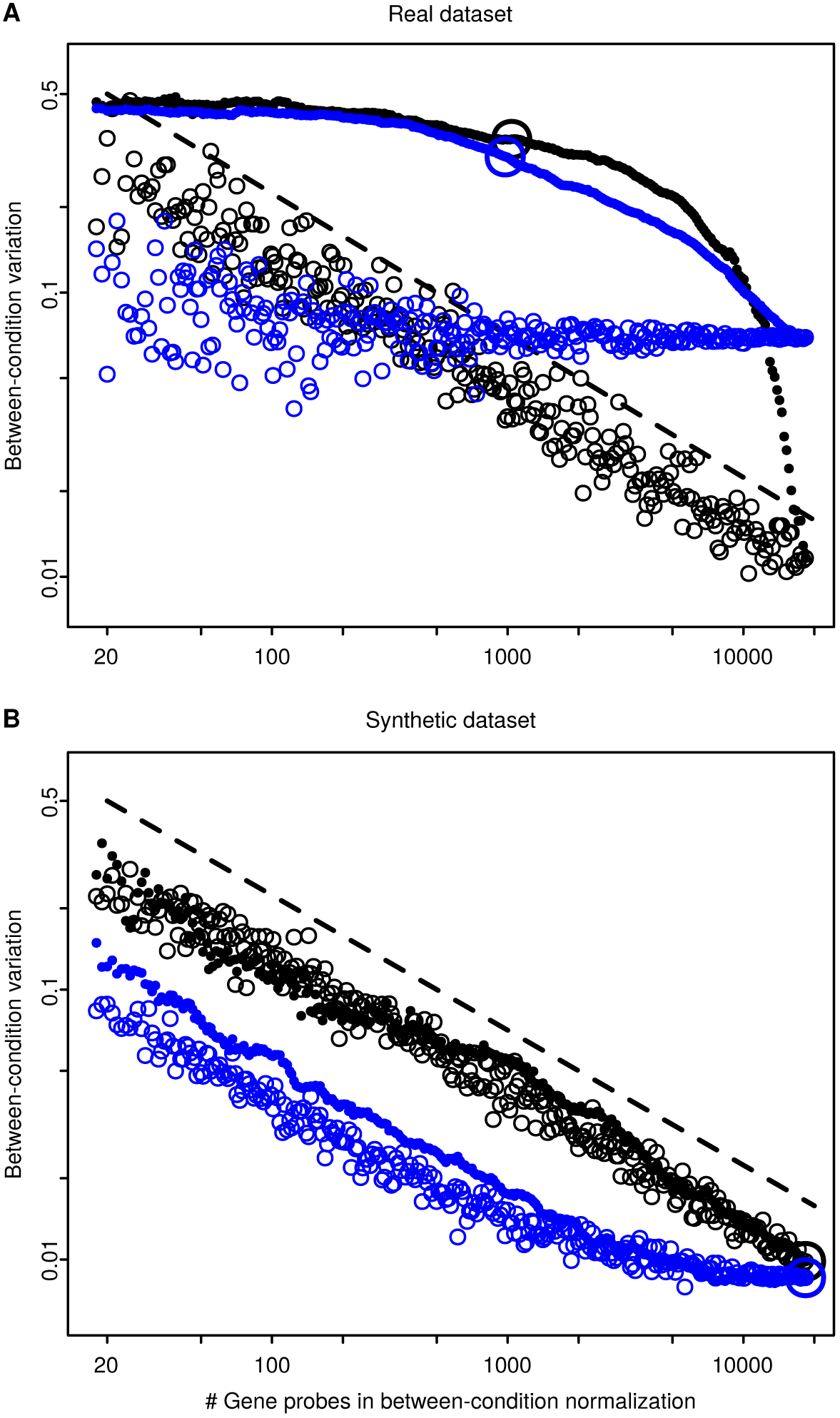
Representing between-condition variation as the standard deviation of the within-condition median averages (averages of sample median expression levels, for all samples under the condition) yields similar results to those obtained with within-condition mean averages (Fig. 2). The panels show the detected variation as a function of the number of gene probes used in the between-condition normalization of the real dataset (A) and synthetic dataset (B). Labeling is the same as in . Each point in either of the panels indicates the variation obtained with one complete normalization (black circles, median condition-decomposition normalization; blue circles, standard-vector condition-decomposition normalization). Gene probes were selected in two ways: randomly (empty circles) or in decreasing order of *p*-values (filled circles). Big circles show the working points of the algorithms whose results are depicted in Figures 1G–J. Black dashed lines show references for *n*decays, with the same values in both panels.

**Supplementary Figure 2:**
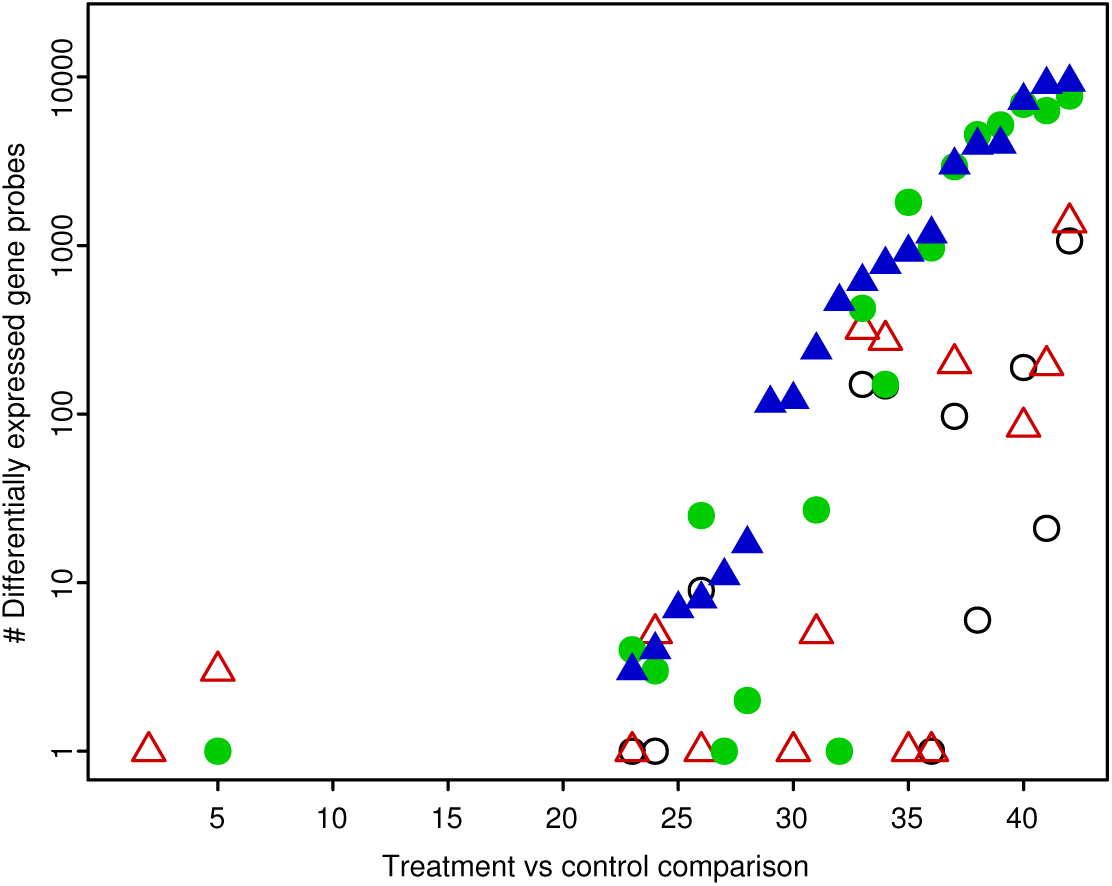
With t-tests, the condition-decomposition normalizations also detected much larger numbers of differentially expressed gene probes (DEGP). The figure shows the number of DEGP obtained with a statistical analysis based on t-tests instead of limma (Fig. 3A). Labeling is the same as in A (empty black circles, median normalization; empty red triangles, quantile normalization; filled green circles, median condition-decomposition normalization; filled blue triangles, standard-vector condition decomposition normalization). Treatment vs control comparisons are ordered according to the number of DEGP identified with standard-vector condition-decomposition normalization, increasing from left to right. This order (not shown) was similar but not exactly the same as in A.

## Supplementary Mathematical Methods

### SM1 Vectorial representation of sample data

Let *x*_1_*,...,x_n_* be the samples of *n* independent and identically distributed random variables *X*_1_*,...,Xn*. Let us represent the samples *x*_1_*,...,x_n_* with the ℝ*^n^* column vector **x** = (*x*_1_*,...,x_n_*)′, and let us denote the sample mean by 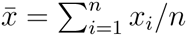.

Let us define the ℝ*^n^* → ℝ*^n^* vectorial operators mean 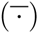 and residual 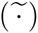, respectively, as

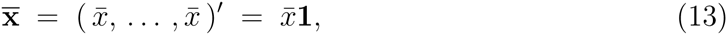

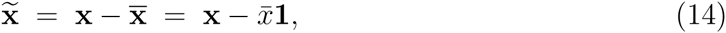

**1** being the all-ones column vector of dimension *n*.

Thus, any sample vector **x** ∈ ℝ*^n^* can be decomposed as

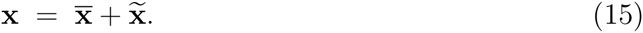

The mean vector 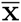 contains the sample mean, while the residual vector 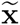 carries the sample variation around the mean.

The vectorial operators mean (13) and residual (14) are linear.

*Proposition*. For any two sample vectors **x, y** ∈ ℝ*^n^* and any two numbers *α, β* ∈ ℝ,

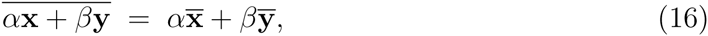

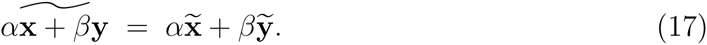

*Proof*. Let us denote **x** = (*x*_1_*,...,x_n_*)′ and **y** = (*y*_1_*,...,y_n_*)′.

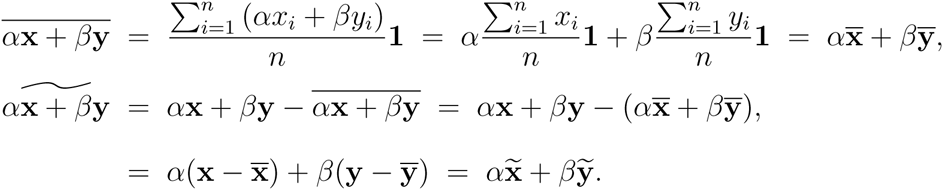

An essential property of the mean and residual vectors is that they belong to subspaces that are orthogonal complements (Eaton, 2007). Hence, for any sample vector **x** ∈ ℝ^n^, the mean vector 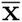 belongs to the subspace of dimension 1 spanned by the unit vector 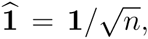, while the residual vector 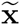 abelongs to the (*n* − 1)-dimensional hyperplane orthogonal to 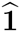.

The lengths of the mean vector and residual vector are equal, up to a scaling factor, to the sample mean and sample standard deviation, respectively. For a set of samples x_1_*,...,xn*, where *n* ≥ 2, let us denote the sample mean as before by 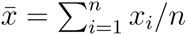, and the sample variance as 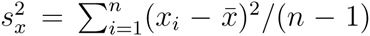. Then, the lengths of the mean and residual vectors obtained from the sample vector **x** = (*x*_1_*,...,xn*)′ are

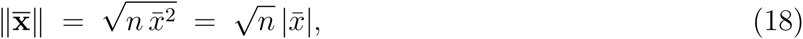

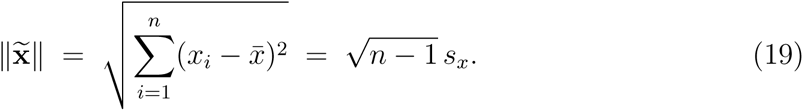

Finally, let us define the standard vector of the sample vector **x** = (*x*_1_*,...,x_n_*)′ (*n* ≥ 2), as

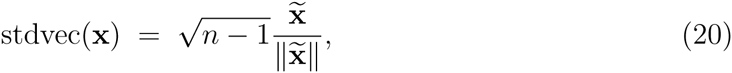

whenever 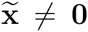, or otherwise as stdvec(**x**) = 0. 0 is the all-zeros column vector of dimension *n*.

For a given number of samples *n*, all the non-zero standard vectors belong to the (*n* − 2)-sphere of radius 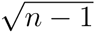, embedded in the (*n* − 1)-dimensional hyperplane perpendicular to 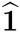. Besides, all the components of a standard vector are equal to the corresponding standardized samples,

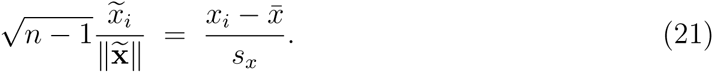

For the degenerate case of having only two samples (*n* = 2), the only possible values of a non-zero standard vector are 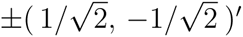′.

### SM2 Linear decomposition of the normalization problem

Let us consider a gene expression dataset, with *g* genes and *c* experimental conditions. Each condition *k* has *s_k_* samples. The total number of samples is 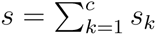.

Let us denote the *observed* expression level of gene *j* in the sample *i* of condition *k* by 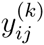. We assume that the observed level 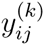 is equal, in the usual log_2_-scale, to the addition of the normalization factor 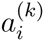 to the *true* gene expression level 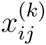,

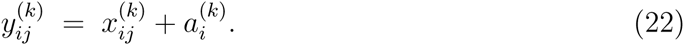

Solving the *normalization problem* amounts to finding the normalization factors 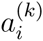 from the observed values 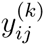. The normalization factors can be understood as sample-wide changes in the concentration of mRNA molecules by multiplicative factors equal to 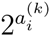. These changes are caused by technical reasons in the assay and are independent of the biological variation in the true levels 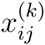.

Let us represent the true and observed expression levels, 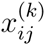 and 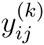, of gene *j* in the samples *i* = 1::: *s_k_* of condition *k*, by the *s_k_*-dimensional vectors

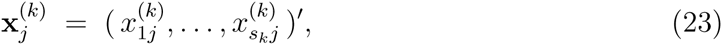

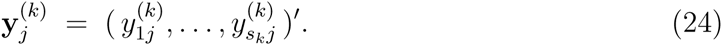

Let us also represent the unknown normalization factors of condition *k b*y the *s_k_*-dimensional vector

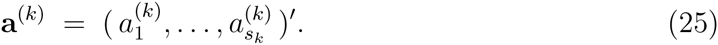

From (22)–(25), the normalization problem can be written in vectorial form as

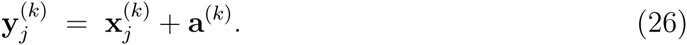

Applying the vectorial operators mean (13) and residual (14), we obtain

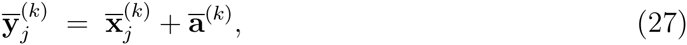

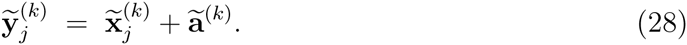

The residual-vector equations (28) correspond to the *c* within-condition normalizations. Each within-condition normalization uses the equations (28) particular to a condition *k*, for the subset of genes 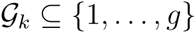 that have expression level available and of enough quality in that experimental condition.

Let us denote the condition means for each gene as

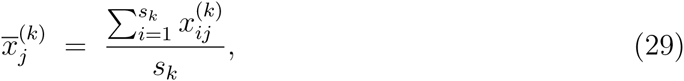

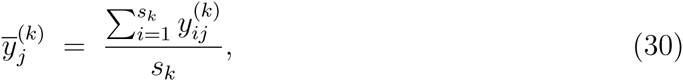

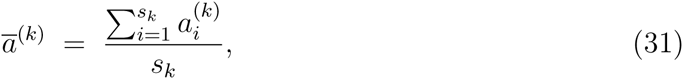

so that

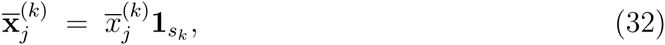

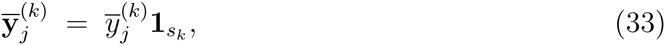

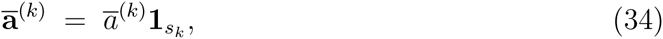

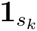 being the all-ones column vector of dimension *s_k_*.

Then, the mean-vector equations (27) can be written as

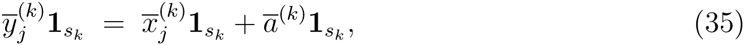

so they reduce to the scalar equations

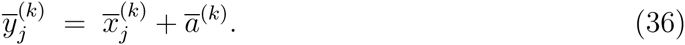

Let us define the vectors of conditions means as

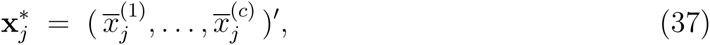

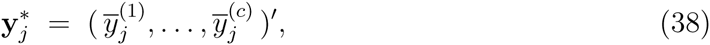

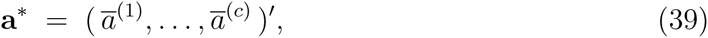

and let us express the condition-mean equations in vectorial form as

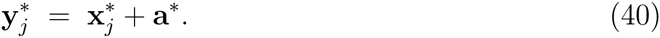

Applying again the mean and variance operators, we obtain

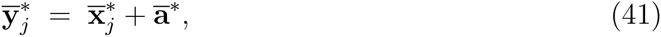

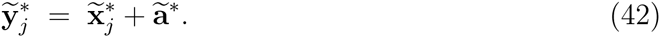

The residual-vector equations on condition means (42) correspond to the single between-condition normalization, in a similar way as (28) do for the each of the within-condition normalizations. There is one equation (42) per gene. The only equations used in the between-condition normalization are those of the subset of genes 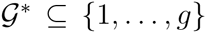 that show no evidence of variation across experimental conditions, according to a statistical test.

Given that 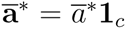, (41) has the only unknown 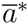. The meaning of 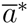 is a conversion factor between the scale the true and observed expression levels. This factor depends on the technology used to measure the expression levels and finding it is out of the scope of the normalization problem. Therefore, without loss of generality, we assume 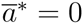, so

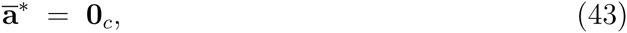

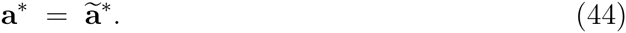

The solution of the between-condition normalization, 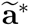, allows to find the mean vectors of the normalization factors 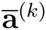, via (34), (39) and (44). The within-condition normalizations yield the residual vectors 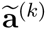. The complete solution to the normalization problem is finally obtained, for each condition *k*, with

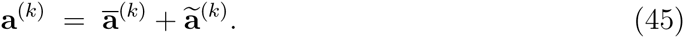

Thus, the original normalization problem (26) has been divided in *c*+1 normalization subproblems on residual vectors, stated by (28) and (42). In fact, this linear decomposition is possible for any partition of the set of *s* samples. The choice of the partition as the one defined by the experimental conditions is motivated by the need to control the biological variation among the genes used in each normalization. All the *c* + 1 normalizations face the same kind of *normalization of residuals problem*, which we define in general as follows.

**Normalization of Residuals Problem**. Let *y_ij_* be the *i*-th observed value of feature *j*, in a dataset with *n* ≥ 2 observations for each of the *m* features. The observed values *y_ij_* are equal to the true values *x_ij_* plus the normalization factors *a_i_*, which are constant across features. In vectorial form, there are *m* equations

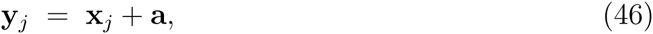
where the vectors belong to ℝ*^n^* . As a consequence

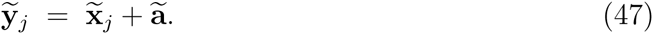

Solving the normalizatio*n of residuals problem amount*s to finding the residual vector of normalization factors 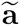 from the observed residual vectors 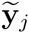. In the within-condition of the corresponding experimental condition. In the between-condition normalization, the features are means of gene expression levels, with one observation per condition.

There is, however, an additional requirement imposed by the methods with which we propose to solve the between-condition normalization. We would like to consider the condition means 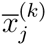 in (36) as sample data across conditions. This only holds when all the conditions have the same number of samples. Otherwise, we balance the condition means so that they result from the same number of samples in all conditions, according to the procedure described in the following.

Let *s*^∗^ be the minimum number of samples across conditions, *s* ^∗^ = min{*s*_1_*,...,s_c_*}. Let 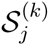 be independent random samples (without replacement) of size *s*^∗^ from the set of indexes {1*,...,s_k_*}, with one sample per gene *j* and condition *k*. Then, the balanced condition means are defined as

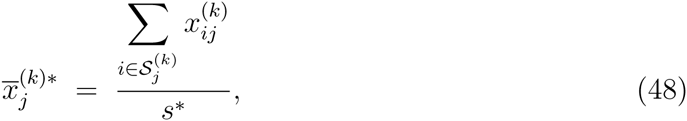

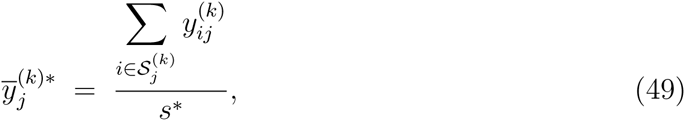

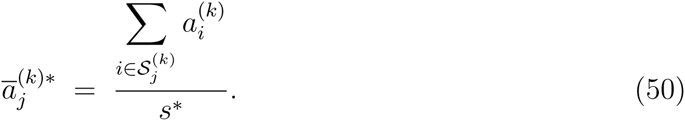

From (22), the balanced condition means verify a relationship similar to (36),

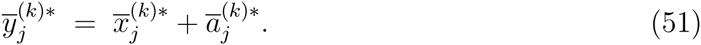

Moreover, the average of 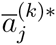 across the sampling subsets 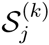 is equal to the unknown 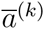. This implies that (51) are, on average, equivalent to (36). Hence, we use the following vectors of balanced conditions means

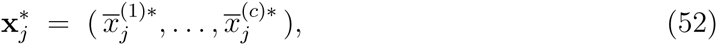

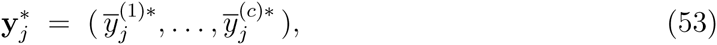

instead of (37), (38), in order to build the condition-mean equations (40). This balancing of the condition means is only required when the experimental conditions have different number of samples.

### SM3 Permutation invariance of multivariate data

Let *x_ij_* and *y_ij_* be, respectively, the true and observed values of a dataset with *n* observations of *m* features, as defined in the *normalization of residuals problem* above.

We have assumed that the *n* true values *x*_1_*_j_,...,x_nj_* of feature *j* are samples of independent and identically distributed random variables X_1_*_j_,...,X_nj_*. These random variables can be represented with the random vector X*j* = (*X*_1_*_j_,...,X_nj_*)′, carried by the probability space (Ω, 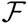, P) and with induced space (ℝ^n^ , 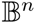 , ℙ). Let us define the random vectors 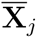 and 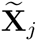 with the vectorial operators mean (13) and residual (14), respectively,

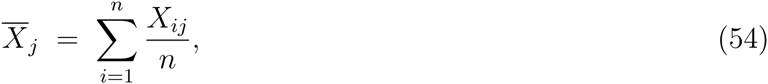

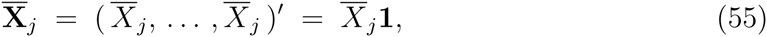

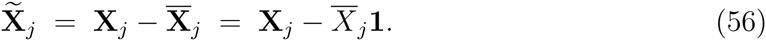

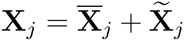 holds for any random vector **X***_j_*, as well as the other properties presented above. Let us assume that E( ||**X***_j_*||) < ∞ and that 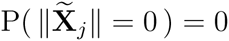, which imply that 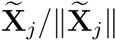 has length 1 almost surely.

The standard random vector 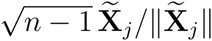 is a pivotal quantity, where the location (mean) and scale (standard deviation) of feature *j* have been removed. The probability distribution of 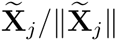 across the remaining degrees of freedom over the unit (*n* − 2)-sphere is governed by the parametric family of the random variables X_1_*_j_,...,Xn_j_*. Moreover, the independence and identity of distribution across the *n* observations implies that the distribution of **X***_j_* is *exchangeable*, i.e. invariant with respect to permutations of the observation labels. As a result, 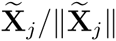 is also permutation invariant, which geometrically corresponds to symmetries with respect to the *n*! permutations of the axes in the *n*-dimensional space of random vectors, projected onto the (*n* − 1)-dimensional hyperplane of residual vectors.

Residual vectors and standard vectors have been widely studied, especially in relation to elliptically symmetric distributions and linear models (Fang et al., 1990; Gupta et al., 2013), and to the invariances of probability distributions (Kallenberg, 2005). Here, we consider these vectors from the viewpoint of the problem of normalizing multivariate data, and its relationship with permutation invariance.

It is well know that, for a multivariate distribution with independent and identically distributed components, the expected value of the standard vector is zero (Eaton, 2007), given that it is so for each component. We prove this here for completeness, and to show that it is also a necessary consequence of the permutation invariance of the distribution.

*Proposition*. The expected value of any true (i.e. without normalization issues) standard vector is zero. If the *n* ≥ 2 samples of feature *j* are independent and identically distributed, then

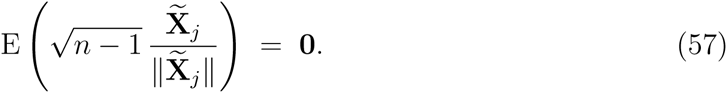

*Proof*. Let 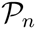 be the set of all the permutation matrices in ℝ*^n^*^×^*^n^*. Then, for any 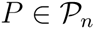, 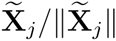 is equal in distribution to 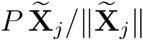. This implies that

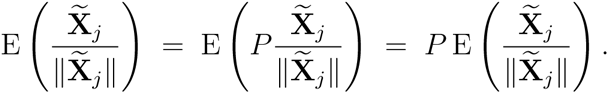

The only vectors that are invariant with respect to all possible permutations are those that have all components identical. Therefore, 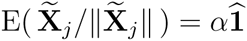, with *α* ∈ ℝ. However, 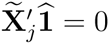, so that 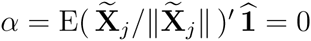. Hence 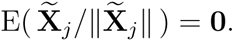

For each true random vector **X***_j_*, there is an observed random vector Y*_j_* = X*_j_* + A, where **A i**s the random vector of normalization factors. The random vectors X*_j_* and **A ar**e independent, representing biological and technical variation, respectively. Therefore, and without loss of generality, we assume in what follows a fixed vector of normalization factors a, i.e. we condition on the event {**A** = **a }**. We also assume that 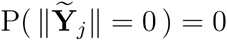, which implies that 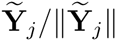 has length 1 almost surely.

In contrast to the true standard vector 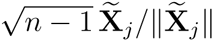, the observed standard vector 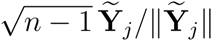 is biased toward the direction of 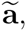 with the result that the expected value is not zero.

*Proposition*. If the *n* ≥ 2 samples of feature *j* are independent and identically distributed, whenever 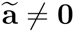,

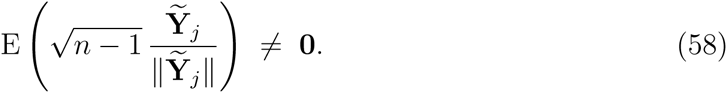

When *n* = 2, there is the additional requirement that 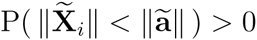. This threshold of detection only occurs for the degenerate case of *n* = 2.

*Proof*. Let us consider the projection of 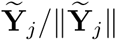 on 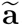, compared to the projection of 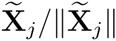.

When the vectors 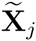 and 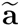 are collinear,

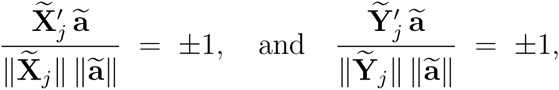

with

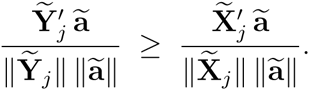

This is the only case when *n* = 2. The additional requirement ensures that, for *n* = 2,

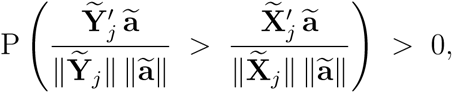

which implies

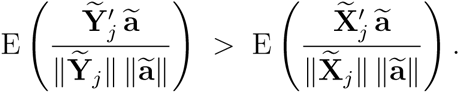

Otherwise, when *n>* 2 and the vectors 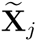 and 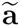 are not collinear, they lie on a plane. The vector 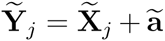 is the diagonal of the parallelogram defined by 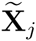 and 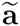. Hence the angle between 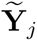 and 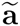 is strictly less than the angle between 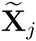 and 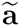, so the cosine of the angle is strictly greater. Thus,

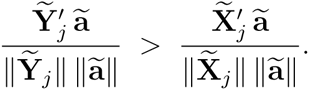

Due to the permutation symmetries in the distribution of 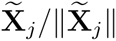, when *n >* 2 the vector 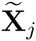 has non-zero probability of being not collinear with 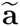, i.e. 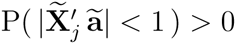.

Therefore,

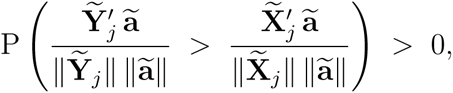

which again implies

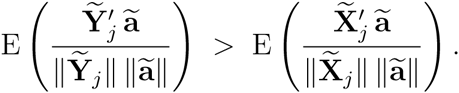

Finally,

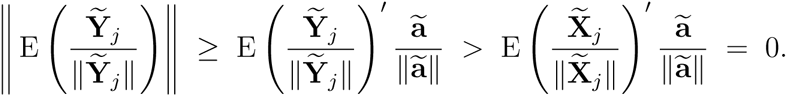

As a consequence, the *normalization of residuals problem* may be restated as the problem of finding the normalization factors 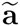 from the observed vectors 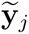, such that the standard vectors 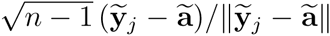 are invariant against permutations of the observation labels. Or equivalently, such that the standard vectors 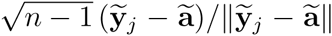have zero mean. The following property provides an approach to the solution.

*Proposition*. Whenever 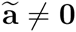, the component of the expected value of 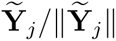 parallel to 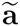 verifies

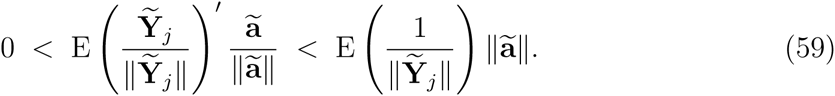

As in (58), when *n* = 2 we also assume that 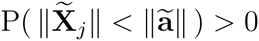.

*Proof*. The first inequality holds from the previous proof. Concerning the second inequality, let us consider

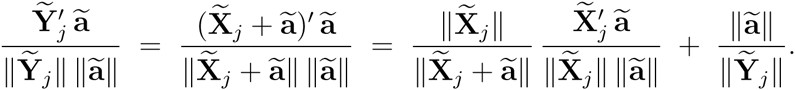

We need to prove that the first term on the RHS has negative expected value. Let us decompose this term into the positive and negative parts,

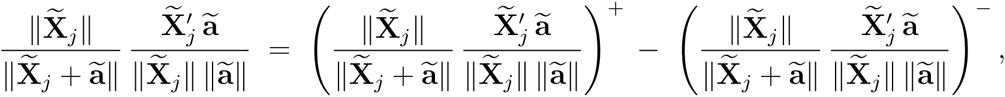
where *X*^+^ = max(*X,* 0) and *X*^−^ = − min(*X,* 0).

Because 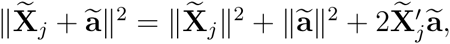

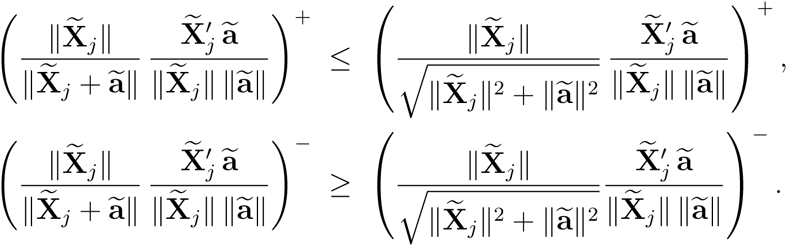

These inequalities are identities when 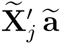 is of opposite sign to ( ·)^±^, or when 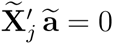. Because of the permutation symmetries of 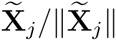, it follows that 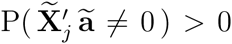, which implies

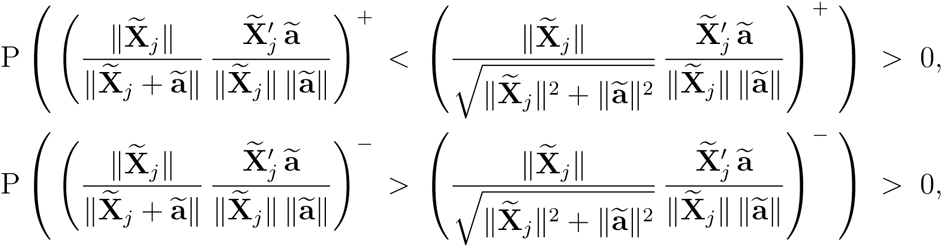

and hence

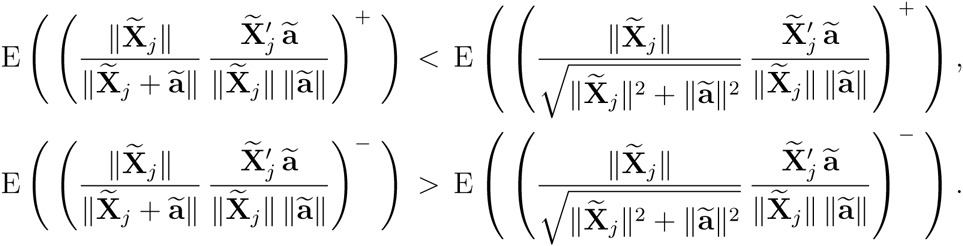

For any permutation matrix 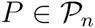,

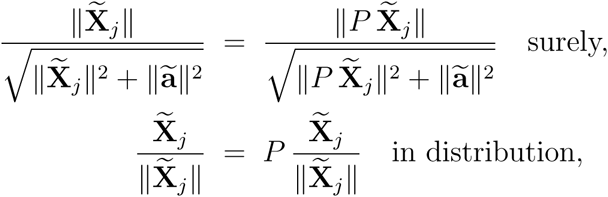

so that

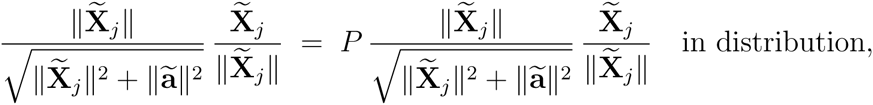

which together with

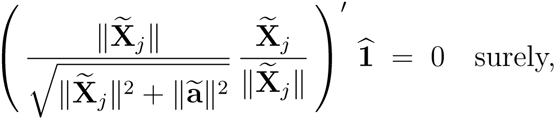

implies, as in (57), that

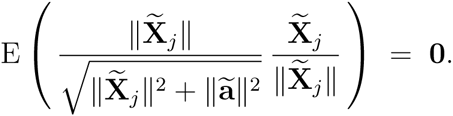

Therefore,

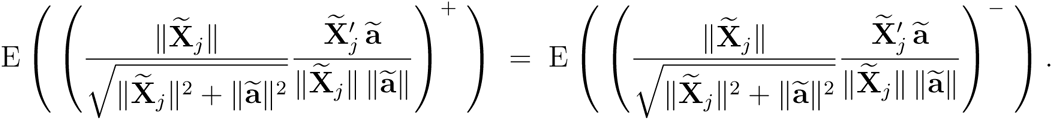

Back to the initial expected values, it follows that

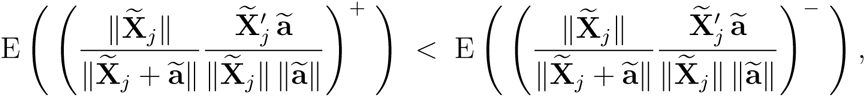

which implies

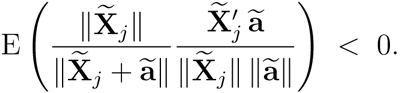

The Gaussian multivariate distribution, among others, has spherical symmetry besides permutation symmetry. For parametric families with spherical symmetry, the true standard vector 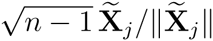 has uniform distribution over the (*n*−2)-sphere. As a result, the components of 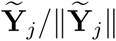 perpendicular to 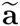 are antisymmetric with respect to the direction of 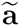, so that they cancel out in expectation. That is, for parametric families with spherical symmetry, and as long as 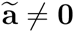,

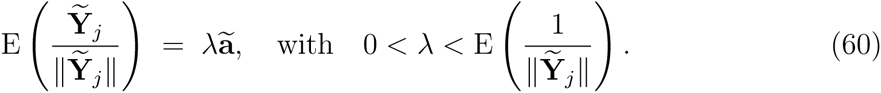

### SM4 Standard-vector normalization

The properties (59), (60) suggest the use of

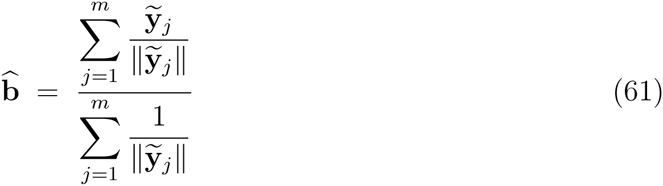

to approximate the unknown residual vector of normalization factors 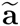. The following iterative method implements this approach to solve the *normalization of residuals problem*.

Let us define the following recursive sequence, where each step *t* comprises *m* vectors 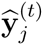 (*j* ∈{1*,...,m*}) and one vector 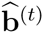,

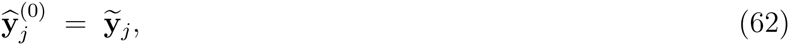

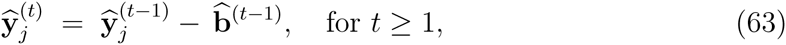

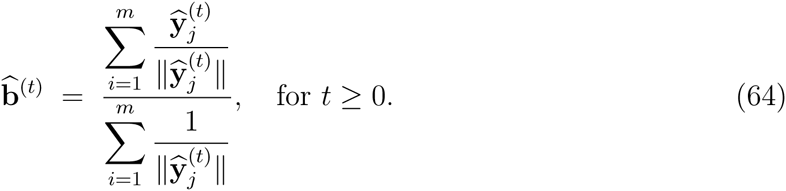

We assume that 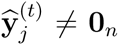, for all *j* ∈{1*,...,m*} and all *t* ≥ 0. Nonetheless, an implementation of this algorithm benefits from trimming out a small fraction (e.g. 1%) of the features with lesser 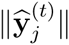 in (64), in order to avoid numerical singularities.

Let us write 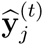 as a function of the unknowns 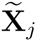 and 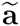. For any *t* ≥ 1,

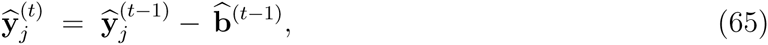

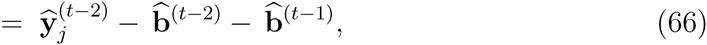

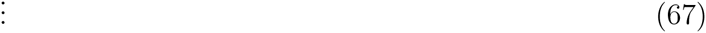

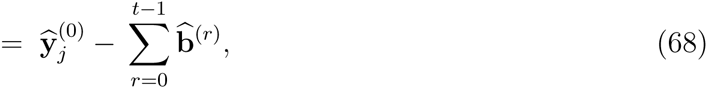

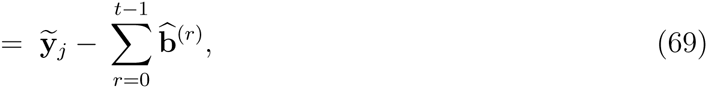

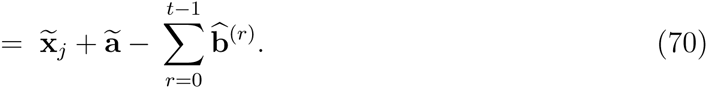

Note that (70) is also valid for *t* = 0.

Let us also define the vectors 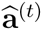, for *t* ≥ 0, which describe the vector of normalization factors still to be removed at step *t*,

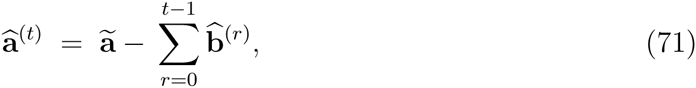

so that, by (70), for *t* ≥ 0,

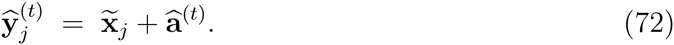

Therefore, the recursive sequence (62)–(64) faces a new, weaker *normalization of residuals problem* at each step *t*, with true residual vectors 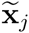, observed residual vectors 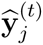 and unknown normalization factors 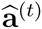. The step *t* results in the estimation of normalization factors 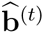, which are removed from 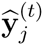, generating the next step. At the beginning, 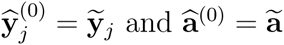.

At convergence, 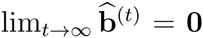. Equations (57), (58), (64) imply that, in such a case, 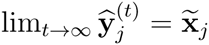 and 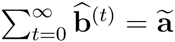. Convergence is optimal when the parametric family of the *m* features has spherical symmetry, Gaussian being the most prominent case. Otherwise, the more uniform the distribution of standard vectors 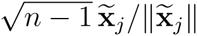 is on the (*n* − 2)-sphere, the faster the sequence (62)–(64) converges. See examples of convergence in Supplementary Movies 1–3.

### SM5 Identification of non-differentially expressed genes

Let us consider a gene expression dataset, with *g* genes and *c* experimental conditions. Each condition *k* has *s_k_* samples. The total number of samples is 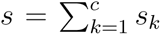. Let us assume that *c* ≥ 2 and that *s_k_* ≥ 2, for all conditions *k* ∈{1*,...,c*}. Let us also assume that, among the *g genes*, there is a fraction *π*_0_ of non-differentially expressed (non-DE) genes, with 0 ≤ *π*_0_ ≤ 1, while the remaining fraction 1 − *π*_0_ comprises the differentially expressed (DE) genes (Storey and Tibshirani, 2003).

Let us consider the usual ANOVA test comparing average expression levels across conditions, gene-by-gene. Under the null hypothesis of a non-differentially expressed gene, the corresponding *F*-statistic follows the *F*-distribution with *c* − 1 and *s* − *c* degrees of freedom. The test of this hypothesis yields a *p*-value *p_j_* for each gene *j* ∈{1*,...,g*}. The obtained *p*-values *p_j_* follow a probability distribution that can be considered as the mixture of two probability distributions, *F*_0_ and *F*_1_, for the non-DE genes and the DE genes, respectively (Storey, 2003). The fraction *π*_0_ of non-DE genes follows the uniform distribution on the interval [0, 1], 

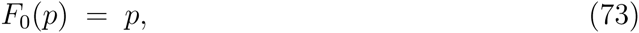
while the fraction 1 −*π*_0_ of DE genes follows a distribution that verifies, for any *p* ∈ (0, 1), 

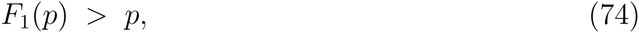
and the mixture distribution is

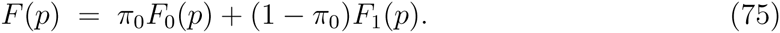

Let us further assume that there exists a *p* ^∗^, with 0 < *p* ^∗^ < 1, such that *F*_1_(*p*) = 1 for every *p* ≥ *p* ^∗^. In other words, all DE genes have *p*-value *p_j_* from the ANOVA test such that *p_j_* ≤ *p* ^∗^, while only some genes among the non-DE genes have p-value with *p_j_ >p* ^∗^. This implies that the mixture distribution of *p*-values is uniform on the interval [*p* ^∗^ , 1],

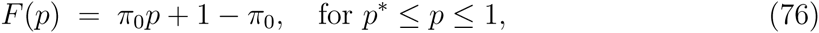

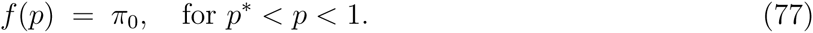

On the other hand, for any set of *n* samples *x*_(1)_ ≤ *x*_(2)_ ≤. . . ≤ *x*_(_*_n_*_)_ obtained from *n* independent and identically distributed uniform random variables on the interval [*a, b*], all the distances between consecutive ordered samples (including boundaries), *x*_(1)_ − *a, x*_(2)_ − *x*_(1)_*,...,x*_(_*_n_*_)_ − *x*_(_*_n_*_−1)_*, b* − *x*_(_*_n_*_)_, obey the same distribution (Feller, 1971). Then, it can be realized that, for any *j suc*h that 2 ≤ *j* ≤ *n* − 1, the two subsets of samples x_(1)_*,...,x*_(_*_j_*_−1)_ and *x*_(_*_j_*_+1)_*,...,x*_(_*_n_*_)_ follow uniform distributions on the intervals [*a, x*_(_*_j_*_)_] and [*x*_(_*_j_*_)_*,b*], respectively.

Based on these facts, to identify non-DE genes we propose finding the minimum *p*_(_*_j_*_)_, from the ordered sequence of *p*-values *p*_(1)_ ≤ *p*_(2)_ ≤ . . . ≤ *p*_(_*_g_*_)_, such that a goodness-of-fit test for the uniform distribution on the interval [*p*_(_*_j_*_)_, 1], performed on *p*_(_*_j_*_+1)_*,...,p*_(_*_g_*_)_, is not rejected. As a result, the genes corresponding to the *p*-values *p*_(_*_j_*_)_*,p*_(_*_j_*_+1)_*,...,p*_(_*_g_*_)_ are considered as non-DE genes.

Given the concavity of *F*(*p*), the goodness-of-fit test used is the one-sided Kolmogorov-Smirnov test on positive deviations of the empirical distribution function.

See Supplementary Movies 4–5 for examples of this approach to identifying non-differentially expressed genes

